# Phylotype-Level Characterization of Complex Lactobacilli Communities Using a High-Throughput, High-Resolution Phenylalanyl-tRNA Synthetase *(pheS)* Gene Amplicon Sequencing Approach

**DOI:** 10.1101/2020.09.09.290726

**Authors:** Shaktheeshwari Silvaraju, Nandita Menon, Huan Fan, Kevin Lim, Sandra Kittelmann

## Abstract

The ‘lactobacilli’ to date encompass more than 270 closely related species that were recently re-classified into 26 genera. Because of their relevance to industry, there is a need to distinguish between closely related, yet metabolically and regulatory distinct species, e.g., during monitoring of biotechnological processes or screening of samples of unknown composition. Current available methods, such as shotgun metagenomics or rRNA-based amplicon sequencing have significant limitations (high cost, low resolution, etc.). Here, we generated a lactobacilli phylogeny based on phenylalanyl-tRNA synthetase (*pheS*) genes and, from it, developed a high-resolution taxonomic framework which allows for comprehensive and confident characterization of lactobacilli community diversity and structure at the species-level. This framework is based on a total of 445 *pheS* gene sequences, including sequences of 277 validly described species and subspecies (out of a total of 283, coverage of 98%). It allows differentiation between 263 lactobacilli species-level clades out of a total of 273 validly described species (including the proposed species *L. timonensis*) and a further two subspecies. The methodology was validated through next-generation sequencing of mock communities. At a sequencing depth of ∼30,000 sequences, the minimum level of detection was approximately 0.02 pg per μl DNA (equalling approximately 10 genome copies per µl template DNA). The *pheS* approach along with parallel sequencing of partial 16S rRNA genes revealed a considerable lactobacilli diversity and distinct community structures across a broad range of samples from different environmental niches. This novel complementary approach may be applicable to industry and academia alike.

**IMPORTANCE:** Species within the former genera *Lactobacillus* and *Pediococcus* have been studied extensively at the genomic level. To accommodate for their exceptional functional diversity, the over 270 species were recently re-classified into 26 distinct genera. Despite their relevance to both academia and industry, methods that allow detailed exploration of their ecology are still limited by low resolution, high cost or copy number variations. The approach described here makes use of a single copy marker gene which outperforms other markers with regards to species-level resolution and availability of reference sequences (98% coverage). The tool was validated against a mock community and used to address lactobacilli diversity and community structure in various environmental matrices. Such analyses can now be performed at broader scale to assess and monitor lactobacilli community assembly, structure and function at the species (in some cases even at sub-species) level across a wide range of academic and commercial applications.

## INTRODUCTION

The generic term ‘lactobacilli’ encompasses all organisms described as belonging to the family Lactobacillaceae until 2020, a highly diverse range of bacteria within the phylum Firmicutes (1). Species within this group are Gram-positive, facultative anaerobic or microaerophilic, rod-shaped, non-spore-forming bacteria. Due to their beneficial effects on food during fermentation, lactobacilli play an important role for the food industry and have thus been studied extensively both for their phenotypic and genomic characteristics over the last decades (2). Several species have received generally recognized as safe (GRAS) status for use in human food and animal feed. Besides their naturally high abundances in fermented foods they are frequently observed as part of the healthy microbiomes of animals, humans and plants (2-4). Formerly, this group consisted of only three genera, *Lactobacillus, Paralactobacillus*, and *Pediococcus*. However, these traditionally defined genera alone represent a genetic diversity larger than that of a typical bacterial family (5, 6). A tremendous recent re-classification effort of the lactobacilli resulted in the establishment of 26 genera (encompassing 272 species plus the proposed species *L. timonensis*) and reduction of the original genus *Lactobacillus* to only 38 species around its type species, *Lactobacillus delbrueckii* (1). This refined taxonomic structure will in future facilitate the detection and description of functional properties that are shared within each genus.

In the last two decades, numerous studies aimed at elucidating the bacterial communities involved in the fermentation of specific traditional foods or beverages by using both culture-dependent and culture-independent methods (reviewed by 7, 8). These communities are most often dominated by lactobacilli. While cultivation-based approaches allow isolation and detailed study of individual strains, they are biased towards microorganisms that are culturable under laboratory conditions (9). Molecular techniques for bacterial community structure analysis in fermented food samples, so far, largely relied on amplification and (high throughput) sequencing of 16S rRNA marker genes (recently reviewed by 10). Comparatively few studies have used shotgun metagenomic sequencing (10-12) despite the availability of highly-resolving bioinformatics tools (13), likely because this approach is still rather costly at a sequencing depth that allows for confident species or even strain level taxonomic classification and, instead, is more suited to derive functional information (14).

Furthermore, the accuracy of taxonomic assignment relies on a curated genome database and the availability of at least one representative genome for every species present in the sample. In the case of the lactobacilli, genome-based taxonomies have been published and updated on a regular basis (5, 6, 15-17). However, new species within this group are even more frequently discovered and described (1).

While draft genomes may not be immediately available, deposition of corresponding marker gene sequences in public databases for reference are required for the valid description of new species. The 16S rRNA gene is a widely accepted marker gene to analyze bacterial and archaeal community diversity and structure in many habitats and to understand evolutionary distances between species. Sequence databases and taxonomic frameworks for bacterial and archaeal 16S rRNA genes such as Greengenes and SILVA are highly curated (18-20) and compatible with next-generation sequencing bioinformatics pipelines such as mothur (14, 21) or QIIME/QIIME2 (22, 23). However, at 16S rRNA gene level and even if the almost entire 16S rRNA gene is analyzed (∼1,500 bp), different lactobacilli type species display sequence similarities greater than the accepted species cut-off of 98.7% and up to 100% sequence identity across shorter regions commonly used for next-generation amplicon sequencing protocols (24, 25). In consequence, high throughput 16S rRNA gene amplicon sequencing provides insufficient taxonomic resolution for understanding the composition of lactobacilli communities in complex environmental samples (26, 27). It is reasonable to speculate that studies using partial 16S rRNA genes as a marker so far merely touched the surface of the vast lactic acid bacterial diversity that may exist in host-associated and fermented food-associated matrices and contribute to their potential beneficial effects. If the aim is not to derive evolutionary relationships but to obtain fine-scale species or even strain-level structural information for a particular sub-community, then other marker genes are available that provide greater taxonomic resolution. Recently, *lacS* and *serB* genes were explored for highly specific strain-level monitoring of *Streptococcus thermophilus* during cheese manufacturing (28-30). For species-level characterization of complex lactobacilli communities across a large number of samples from different matrices the use of the internal transcribed spacer (ITS) region (31) or the *groEL* gene (32) has been suggested by different groups. The ITS region has the advantage of highly conserved primer binding sites due to the flanking 16S and 23S rRNA genes. However, difficulties to extract ITS sequence information from publicly available (draft) genomes has hampered the taxonomic assignment of the majority of lactobacilli species, and the use of a multi-copy gene region, such as the ITS presents an additional challenge (31). Moreover, both, ITS and *groEL* do not seem to provide a clear separation between several closely related species (e.g., species belonging to the *L. casei, L. plantarum, L. sakei, L. delbrueckii*, and *L. buchneri* phylogroups as defined earlier (1, 6)).

One of the most well-characterized marker genes of lactobacilli is the phenylalanyl-tRNA synthetase (*pheS*) gene. For the genera *Enterococcus, Lactobacillus* and *Pediococcus*, Naser *et al*. (26, 33) reported a *pheS*-based interspecies gap which, in most cases, exceeded 10% sequence dissimilarity, and an intraspecies variation of up to 3% across an alignment consisting of approximately 450 bp (26, 33). Currently available next-generation sequencing chemistry easily achieves full coverage of amplicons of this size, thus allowing to make full use of the resolving power of the *pheS* gene. These characteristics along with the general practice to deposit the *pheS* gene along with the valid description of novel species makes the *pheS* gene a highly suitable candidate for lactobacilli diversity and composition analysis in fermented foods or other host-or non-host associated environmental samples.

Here, we introduce a new approach for high-throughput next-generation sequencing based on *pheS* marker genes and a curated *pheS*-based taxonomic framework to characterize diversity and structure of lactobacilli communities down to the species level. We demonstrate validity of this method by analysis of mock lactobacilli communities (containing DNA of 13 different lactobacilli (sub)species belonging to 7 different genera, which differed in the known relative amounts of each taxon). The robust taxonomy allowed us to assess lactobacilli community structure in samples from complex host-and food-associated environments at high-throughput and high-resolution.

## MATERIALS AND METHODS

### In silico assessment of variability and coverage of pheS gene primers

For assessment of primer binding site variability and primer coverage, draft genomes of 266 lactobacilli species and subspecies were downloaded from NCBI (Tab. S1 in the supplemental material). Available *pheS* sequences were used to BLAST search these genomes for candidate regions with an e-value of 5 to filter spurious hits.

Subsequently, these regions were extended by 400 bp at either end and excised from the genomes for subsequent primer binding site analysis. The obtained sequences were aligned in MUSCLE (34) and subjected to the WebLogo 3 tool for visualization of sequence variability of the *pheS* gene along the primer binding sites (35). The percentage of primer coverage was calculated from the alignment by establishing the number of mismatches of each type strain with the primer sequences.

### Establishment of pheS gene sequence database and taxonomic framework

A total of 445 *pheS* gene sequences were collected for this study. They were either directly downloaded from NCBI or obtained in this study *via in silico* PCR or *via* BLAST search against whole genome sequences of type strains. These sequences cover the type strain sequences of 267 *Lactobacillus* and *Pediococcus* species and 10 subspecies. Of these, 164 were available from previous publications, however, 3 were replaced with sequences derived by *in silico* PCR of the type strains (this study; Tab. S1 in the supplemental material). Reference sequences from a total of 113 species were previously unavailable and were obtained in this study by using *in silico* PCR (77 sequences, not including that of *L. curieae* 2) or BLAST search (36 sequences) against the genomes of the type strains. Briefly, whole genome assemblies of the type strains were downloaded from NCBI. *In silico* PCR was performed using the Cutadapt software (36) and primers as described earlier (26). Species, for which the *in silico* PCR approach failed, were subjected to BLAST search and extraction of the *pheS* sequence of the closest relative at whole genome level (if known) or at marker gene level by using an in-house Python script.

Sequences of the six remaining type species could not be obtained in this study (Tab. S1 in the supplemental material). All 445 *pheS* gene sequences were imported into the ARB software package (37) and aligned. A phylogenetic tree was constructed using the Neighbor-Joining method including 384 sequences (*pheS* gene alignment positions 487-884) and Jukes-Cantor correction. The remaining 61 sequences were added to the tree using the FastParsimony Tool in ARB (*pheS* gene alignment positions 556-802). Type strain sequences that shared ≥97% sequence similarity were regarded as non-resolvable based on *pheS* and given the same species-level clade name. Non-type strain sequences that shared ≥90% *pheS* gene sequence similarity with the closest type strain were assigned to the same species-level clade. However, sequences are still individually traceable based on their unique strain-level identifiers (=accession numbers). Non-type strain sequences that shared <90% *pheS* gene sequence similarity with the closest type strain, were subjected to further investigations to assess their standing in nomenclature based on whole genome sequence similarity analysis as follows: Draft genomes of the closest type strains were downloaded from NCBI, and average nucleotide identity (ANI) values were calculated using the average_nucleotide_identity.py script from the pyani package (38). We used the typical percentage threshold of 95% for the ANI-based species boundary (39). Non-type strains that shared <90% *pheS* gene sequence similarity and <95% ANI with the closest related type strain, were given the clade name of the closest type strain, but were assigned a separate clade number (*e*.*g*., *L. curieae* 2).

Subsequently, each sequence was assigned from phylum down to species (sub-species where applicable) and strain level (=accession number) using the Greengenes scheme (18) according to the list of prokaryotic names with standing in nomenclature (http://www.bacterio.net/index.html). In addition to sequences belonging to the former genera *Lactobacillus, Paralactobacillus*, and *Pediococcus, pheS* gene sequences of the type species of each of the genera *Enterococcus* (*E. faecalis*, AJ843387), *Fructobacillus* (*F. fructosus*, AM711194), *Lactococcus* (*L. lactis*, JN226442), *Leuconostoc* (*L. mesenteroides*, AM711145), *Oenococcus* (*O. oeni*, FM202093), *Streptococcus* (*S. pyogenes*, AM269572), and *Weissella* (*W. viridescens*, FM202120) as well as the *pheS* gene sequence of *Bacillus subtilis* (extracted by BLAST search against the type strain ATCC 6051; CP003329) were also included in the taxonomic framework.

### Construction of Lactobacilli mock communities

A total of 13 lactobacilli strains were individually grown in MRS media (Tab. S2 in the supplemental material). Nucleic acids were extracted by using bead-beating in combination with the Maxwell DNA extraction system. Briefly, 30 μl of pellet from the overnight culture was weighed into a bead-beating tube (MP Biomedicals LLC, Santa Ana, CA, USA) together with 550 μl buffer A (NaCl 0.2 M, Tris 0.2 M, EDTA 0.02 M; pH 8), 200 μl 20% SDS, and 20 μl Protease K (Promega, Madison, WI USA). The sample was incubated for 30 min at 56°C. Bead-beating was performed at 6.0 m/s for 40 s using the FastPrep24 system (MP Biomedicals LLC). The sample was centrifuged at 16,000 × *g* for 6 min. The supernatant was transferred into the first well of the Maxwell cartridge containing 300 μl of lysis buffer, and all subsequent steps were done as described in the manufacturer’s protocol (Maxwell 16 FFS Nucleic Acid Extraction System, Promega). DNA was eluted into a total volume of 80 μl elution buffer (EB, 10 mM Tris, pH 8.5 with HCl). Subsequently, nucleic acids from individual strains were serial diluted and mixed at known concentrations to generate two different mock communities (Tab. S2 in the supplemental material). The two final pools, each containing a total of 100 ng DNA were evaporated and re-eluted in 50 μl of RNase-free water. Per PCR reaction of 20 μl, 1 μl of the mock mix DNA was used.

### Collection and processing of fermented food and host-associated samples and extraction of nucleic acids

In this study, 24 samples from various environments were analyzed (Tab. 1). Fermented food samples were purchased from local markets or home-fermenting individuals in three countries in South-East Asia: Vietnam, Indonesia, and Malaysia. Coconut milk yogurt, Greek yogurt and kefir, all labelled to contain live cultures were obtained from an organic store in Singapore. Aliquots of a total volume of approximately 20 ml were freeze-dried using a VIRTIS lyophilizer (SP Industries, Gardiner, NY, USA) and subsequently homogenized using a coffee grinder (CoffeeMill, Severin, Germany), which was thoroughly sterilized in-between samples. Nucleic acids were extracted from 30 mg of food-associated sample as described above for lactobacilli isolates by using the combined bead-beating and Maxwell DNA extraction protocol.

To test applicability of the method to host-associated samples, a fresh fecal sample each was collected from a free-ranging wild boar (Singapore, NParks research permit number 18-075) and a pet guinea pig (Singapore). Nucleic acids were extracted with the Qiagen Power Fecal Pro kit (Qiagen, Hilden, Germany) according to the manufacturer’s protocol. In addition, DNA from the jejunum content of a laboratory mouse fed on a standard chow diet was extracted according to Rius *et al*. (40, kindly provided by Henning Seedorf, Temasek Life Sciences Laboratory, Singapore).

### Amplification and next-generation sequencing of 16S rRNA and pheS marker genes

PCR amplification of 16S rRNA and phenylalanyl-tRNA synthetase α-subunit (*pheS*) genes was carried out using previously described primers 515F (5’-GTGYCAGCMGCCGCGGTAA-3’; 41) – 806R (5’-GGACTACNVGGGTWTCTAAT-3’; 42), and pheS21F (5’-CAYCCNGCHCGYGAYATGC-3’; 26) – pheS23R (5’-GGRTGRACCATVCCNGCHCC-3’; 26), respectively, and PCR cycling conditions as described previously (26, 43) on a Bio-Rad T100 thermal cycler (Bio-Rad Laboratories Inc, Hercules, CA, USA). The expected amplicon sizes were approximately 330 bp for the 16S rRNA gene fragment and 431 bp for the *pheS* gene fragment. The PCR mixture in each assay contained 30 µl REDiant 2× PCR Master Mix (1st BASE, Axil Scientific Pte Ltd, Singapore), 6 µl forward primer (final concentration 0.2 µM), 6 µl reverse primer (0.2 µM), 14 µl RNase-free water (Life Technologies, Grand Island NY, USA) and 3 µl of DNA (samples with DNA concentrations above 60 ng per µl were diluted). The assay was split into triplicate reactions per DNA sample. Three reactions per 96-well plate did not contain any DNA and served as negative controls. Successful amplification and absence of amplification products from the no-template negative controls was verified on 1% [w/v] agarose gels. Subsequently, amplicons were sent to NovogeneAIT (Singapore, Singapore) for multiplexing and sequencing on a NovaSeq 6000 sequencer using PE250 chemistry (Illumina, San Diego, CA, USA).

### Bioinformatic and statistical analyses

High throughput 16S rRNA and *pheS* gene amplicon sequence data was processed using the QIIME2 pipeline (23). Paired-end reads were filtered, denoised, merged, and chimera-checked by using DADA2 (44). Parameters for 16S rRNA gene sequences were “--p-trim-left-f 27 --p-trim-left-r 28 --p-trunc-len-f 200 --p-trunc-len-r 150 --p-chimera-method pooled”, while parameters for *pheS* gene sequences were “--p-trim-left-f 27 --p-trim-left-r 28 --p-trunc-len-f 0 --p-trunc-len-r 0 --p-chimera-method pooled”. DADA2-derived representative sequences of bacterial 16S rRNA genes were taxonomically assigned with SILVA (version 138; https://www.arb-silva.de/silva-license-information/, 19) using the QIIME2 compatible files produced according to the Github pipeline (https://github.com/mikerobeson/make_SILVA_db). DADA2-derived representative sequences of *pheS* genes were taxonomically assigned using *pheS*-DB (this study). For 16S rRNA gene sequence data, classification was done using the vsearch algorithm with default parameters (“--p-maxaccepts 10” and “--p-perc-identity 0.80”, 45). For *pheS* gene sequence data, the vsearch algorithm was applied using default parameters apart from “--p-maxaccepts 1” and “--p-perc-identity 0.90” (45). For downstream analyses, all abundance tables were collapsed at genus (-L 6) and species level (-L 7), exported to .tsv format, and further processed in Excel (Microsoft Corp., Redmond, WA, USA).

The fraction of representative sequence reads that was classified as “unassigned” by *pheS*-DB was dissected further using Kraken2 (reference database version 2019, 46). *PheS*-DB-unassigned representative sequence reads that were assigned to the lactobacilli by Kraken2 were BLAST searched against the NCBI Nucleotide collection (nr/nt) for differentiation between true *pheS* and unspecific sequences. True *pheS* hits were imported into ARB, aligned and calculated into the phylogenetic tree for additional verification. The total percentage of sequence reads that these representative sequences accounted for was calculated from the respective feature table in QIIME2.

To compare species richness between samples analyzed based on 16S rRNA and *pheS* genes, Chao-1 indices were calculated using the software PAST (47). The similarity of environmental samples within and between methods was assessed using the Bray-Curtis similarity metric in PAST (47, 48).

Student’s t-test (two-tailed with unequal variance) was performed in Excel (Microsoft) and used to test for significant differences between Chao-1 indices based on 16S rRNA and *pheS* genes.

Visualization of sample similarity by clustering was done by generation of a heatmap and dendrograms in ClustVis, where row centering and unit variance scaling were applied, and both, rows and columns, were clustered using correlation distance and average linkage (49). Principal Component Analysis (PCA) was also performed in ClustVis. For this, unit variance scaling was applied to the rows, and principal components were calculated using singular value decomposition with imputation.

### Data availability

Partial *pheS* gene sequences from isolated lactobacilli were deposited with GenBank under accession numbers MT415913 – MT415925. Partial 16S rRNA and *pheS* gene sequence data generated by high-throughput amplicon sequencing were deposited in the NCBI BioProject database under accession numbers PRJNA629775 (reviewer link: https://dataview.ncbi.nlm.nih.gov/object/PRJNA629775?reviewer=fis4o9g9nckdtnldre8h9262pg).

## RESULTS AND DISCUSSION

### Evaluation of pheS primer binding sites and coverage within the lactobacilli

Here, we assessed the *pheS* gene as an alternative marker gene for high-resolution profiling of lactobacilli. Primer pairs pheS21F/pheS22R and pheS21F/pheS23R had been reported previously to successfully target a wide diversity of *Enterococcus* and former Lactobacillaceae species *in vitro* (26, 33). Since the time of these earlier publications, numerous additional lactobacilli species have been validly described. While the amplicon of primer pair pheS21F/pheS22R is considerably longer (494 bp), primer pair pheS21F/pheS23R results in an amplicon of 431 bp in length, which makes it suitable for high-throughput next generation sequencing with currently available technologies (Fig. 1A, B). In view of the increased lactobacilli diversity that has been unraveled in recent years, we aimed to re-evaluate sequence conservation along the primer binding regions and coverage of the primers amongst the target species. The *pheS* gene region was excised from a total of 266 lactobacilli type strains, for which genomes are publicly available (273 out of 283 type strains including all subspecies and the proposed species *L. timonensis*; for *B. bombi*, the type strain genome was unavailable and that of strain BI-2.5 was used instead).

**FIG 1.**
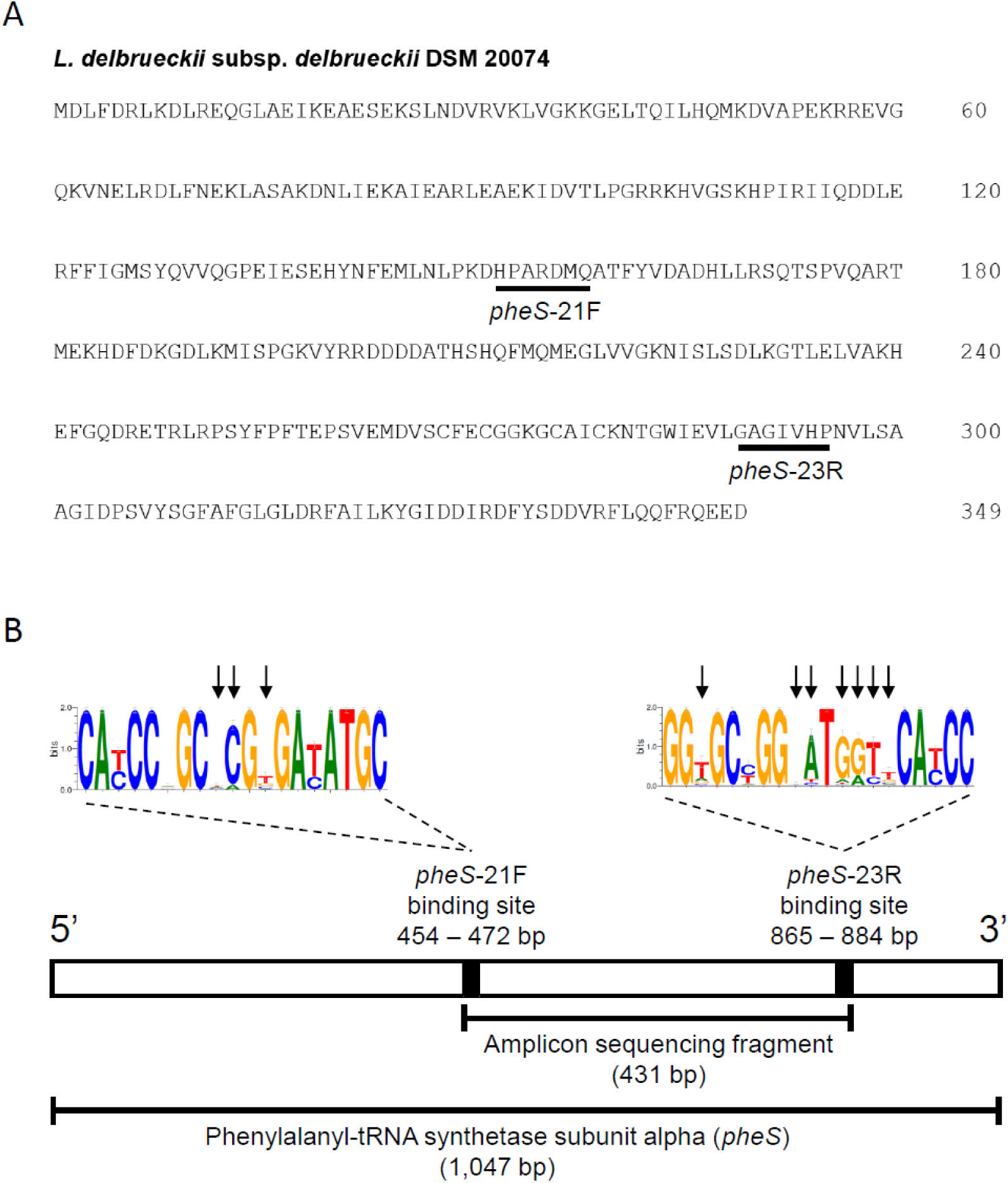
Primary structure of the complete phenylalanyl-tRNA synthetase subunit alpha (*pheS*) and binding sites of the primers used in this study. A. Protein sequence of the *Lactobacillus* type species, *L. delbrueckii* subsp. *delbrueckii* DSM 20074 with amino acid numbering. Approximate binding sites of primers pheS21F and pheS23R are indicated. B. Graphical map of the complete *pheS* nucleotide sequence depicting the positions of primer binding and PCR amplicon length. Primer sequence conservation was tested across 266 lactobacilli type strain sequences and is represented through a sequence web logo. The overall height of each stack indicates the sequence conservation at that position, the height of symbols within each stack indicates the relative frequency of nucleic acids at that position, and error bars indicate an approximate Bayesian 95% credible interval. Black arrows denote positions at which mismatches to the primers were observed.

Extracted *pheS* genes were aligned, and representations of the consensus sequences along the primer binding regions were visualized through sequence web logos (35, Fig. 1B). As expected, the degree of conservation varied between primer binding positions. For pheS21F, none of the tested species showed >2 mismatches with the primer sequence, suggesting comprehensive coverage for the target community. Importantly, primer pheS21F had no mismatches with the crucial five consecutive bases counted from the 3’ end of the primer sequence. A higher degree of sequence variability was observed for the binding region of primer pheS23R (Fig. 1B), and 9% of the sequences showed a mismatch at position 3 from the 3’ end. However, the majority of lactobacilli type strains had ≤2 mismatches with the primer sequence (82%). In future, and making further compromise on specificity, it may be possible and desirable to increase degeneracy of the primer sequences to test if coverage extends.

### pheS gene phylogeny of the lactobacilli

Our comprehensive *pheS* phylogeny encompasses sequences from 277 out of the total 283 lactobacilli species and subspecies validly described by March 2020 (coverage of 98%, Fig. 1, Tab. S1 in the supplemental material). Representative *pheS* gene sequences of the remaining six type species could not be derived in this study, either, because *in silico* PCR and BLAST from publicly available draft genomes were unsuccessful (2 type species) or because the genome of a particular described species has not been sequenced or deposited in public databases yet (4 type species, Tab. S1 in the supplemental material). Our analysis revealed several inconsistencies in the species assignments of previously deposited *pheS* gene sequences. Thus, seven previously published sequences were re-classified based on the comparison of *pheS* gene sequence similarities to the type strain of the previous species assignment and to other closely related type strains (Tab. 2, Fig. 2).

**FIG 2.**
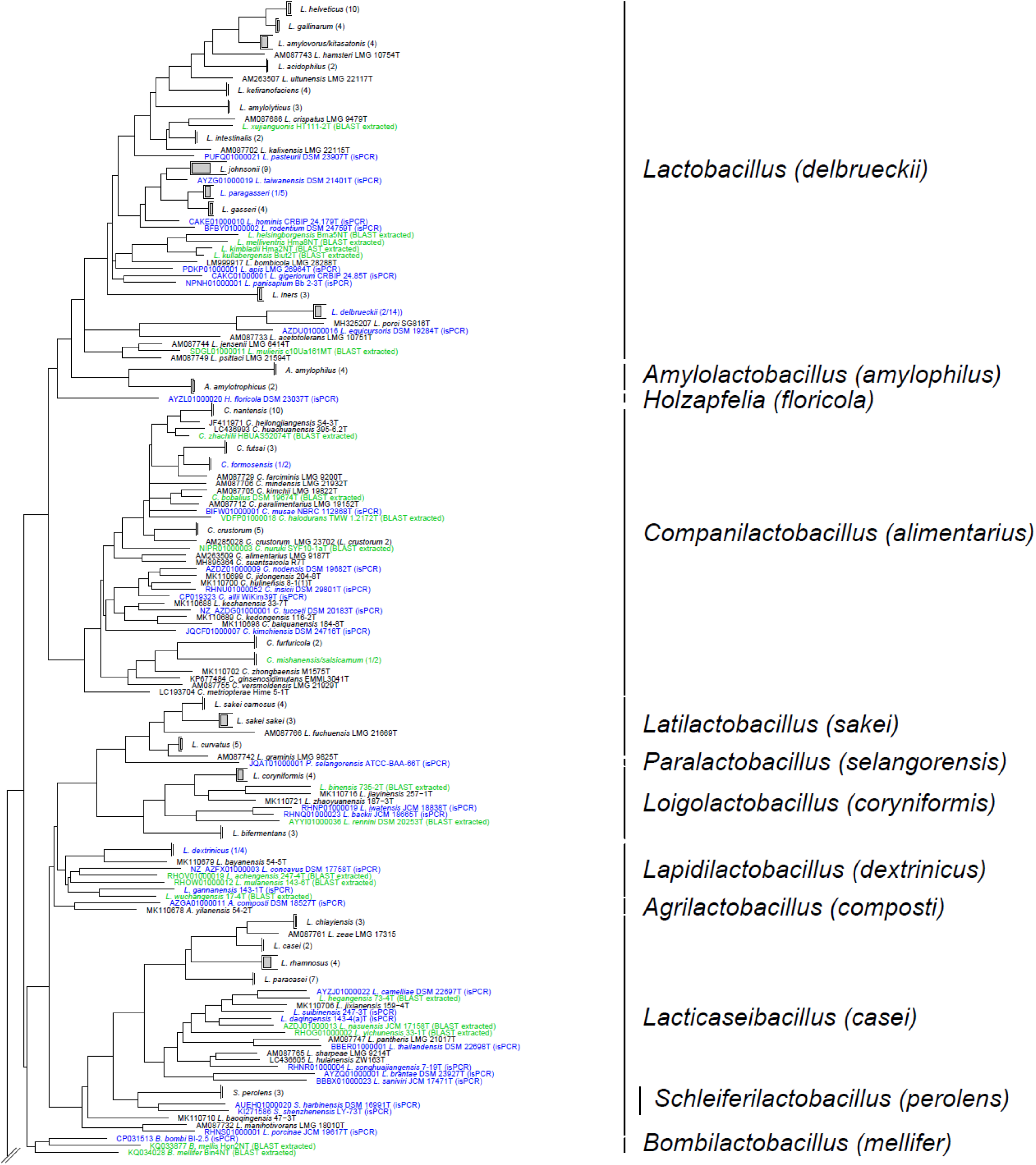
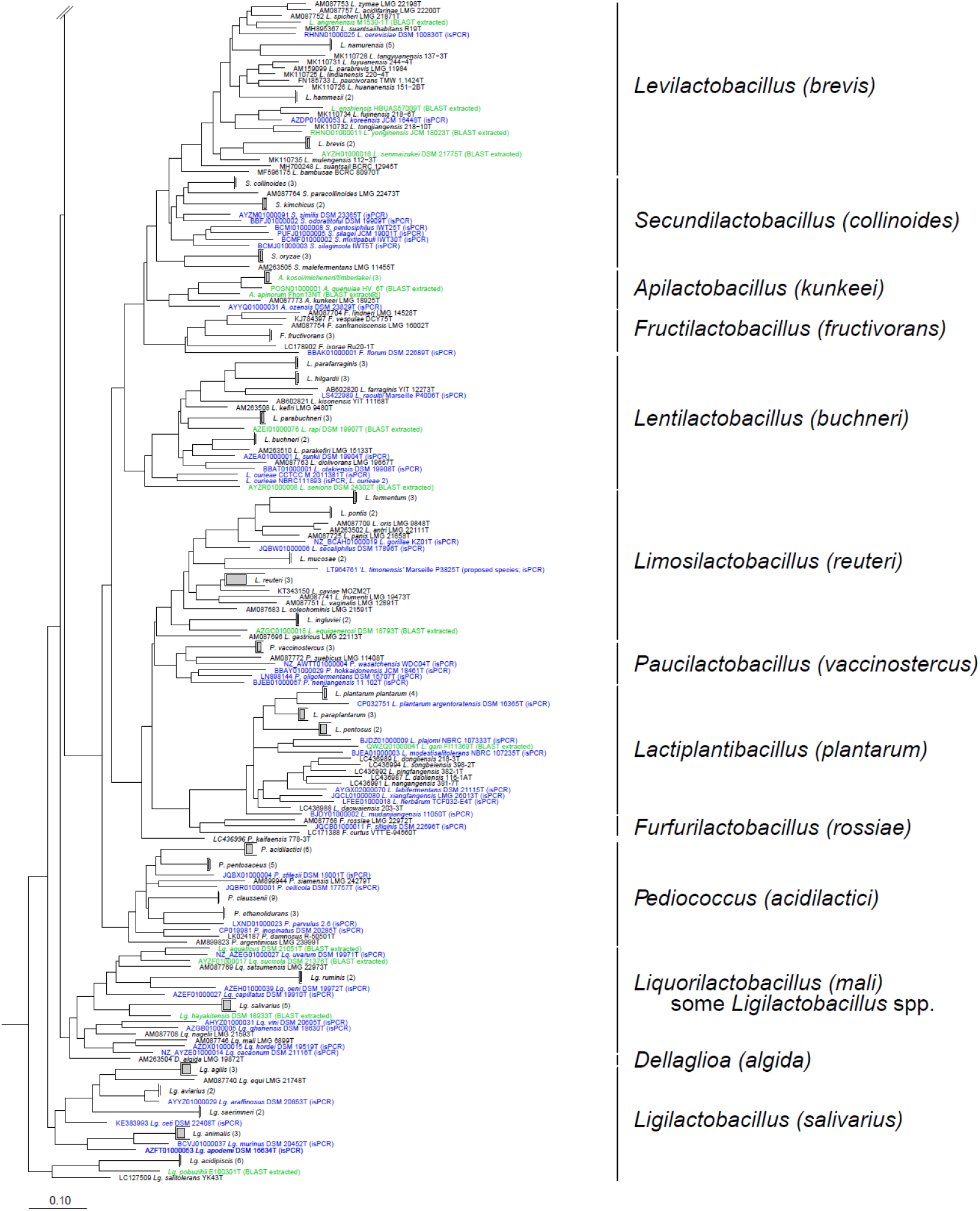
Phylogenetic tree showing 263 species-level clades derived from a total of 445 lactobacilli phenylalanyl-tRNA synthetase (*pheS*) gene sequences. The backbone, consisting of 384 sequences (alignment positions 487-884), was calculated in ARB using the Neighbor-Joining algorithm with Jukes-Cantor correction. The remaining 61 partial sequences (alignment positions 556-802) were added using the Fast Parsimony tool. The scale bar indicates 0.1 nucleotide substitution per nucleotide position. For clarity, some coherent groups of sequences are indicated only as triangles or trapezoids, with the number of sequences clustering into these groups given in parentheses behind the clade name. Sequences in green were derived from isPCR, while those in blue were obtained by BLAST-based extraction from the draft genomes. The *pheS* gene sequences of 16 *Enterococcus faecium* strains (including the type strain LMG 11423^T^; GenBank accession number AJ843428) served as the outgroup. New genus-level affiliations and former *Lactobacillus* and *Pediococcus* phylogroups (in brackets) are indicated to the right of each clade. Generally, the former phylogroup name matches exactly the type species of the new genus. However, in the case of the former “algidus” phylogroup, the type species has been renamed as *Dellaglioa algida*. Genus affiliations of sequences belonging to *Ligilactobacillus* (*Lg*.) and *Liquorilactobacillus* (*Lq*.) are abbreviated with two letters for clarity.

The average observed *pheS* gene inter-species sequence similarity was 69.7% ± 4.9% (± standard deviation), with the inter-species gap of neighboring species exceeding 10% in most cases. The average intra-species sequence similarity was 99.2% ± 1.6% (StDev), which is comparable to the intra-species dissimilarity of up to 3% reported earlier (33). Most type strains shared <97% *pheS* gene sequence similarity. The chosen arbitrary threshold allowed for clear species calling for all but seven validly described species. *L. amylovorus* and *L. kitasatonis* shared 98.5% *pheS* gene sequence similarity, while *Apilactobacillus kosoi, A. micheneri* and *A. timberlakei* shared 98.4–99.5% *pheS* gene sequence similarity (Tab. 3).

*Companilactobacillus mishanensis* and *C. salsicarnum* shared 100% *pheS* gene sequence similarity, and an Average Nucleotide Identity (ANI) of 99.9%. Despite their recognition as separate species, they appear to not be easily differentiated on basis of the *pheS* gene alone. Here, we named the clades “*L. amylovorus/kitasatonis*”, “*A. kosoi/micheneri/timberlakei*”, and “*C. mishanensis/salsicarnum*” (Tab. 3).

Strains formerly recognized as *L. zeae* now belong to the species *Lacticaseibacillus casei* (1). However, these isolates, which appear to be different at the strain level, are confidently distinguished using *pheS*-DB (if selecting strain level resolution “-L 8”). Our *pheS* gene sequence analysis further indicated that strain NBRC 111893, previously deposited as *Lentilactobacillus curieae* NBRC 111893, only shared 84.4% *pheS* gene sequence similarity with the type strain *L. curieae* CCTCC M 2011381^T^ (*pheS*-DB clade name “*L. curieae*”, 50) and an ANI of only 83.8% (Tab. 3). This suggests that strain NBRC 111893 may represent a novel species. Hence, it was given the clade name “*L. curieae* 2”). Similarly, *pheS* gene sequence analysis confirmed a previous observation by Scheirlinck and colleagues that *Companilactobacillus crustorum* strain LMG 23702 is considerably different from the type strain *C. crustorum* LMG 23699^T^ (≤89.9% *pheS* gene sequence similarity; 51). Unfortunately, no genome is available for strain LMG 23702 for analysis at the genomic level. Until more data become available, we allow the *pheS* approach to differentiate between the *C. crustorum* clade that includes the type strain (“*C. crustorum*”) and strain LMG 23702 (“*C. crustorum* 2”). In future, a polyphasic approach should be used to characterize strains NBRC 111893 and LMG 23702 at the genomic and physiological level, thereby providing clarity on their potential roles as the type strains of novel species.

Until recently, seven lactobacilli species were subdivided into subspecies. The former two subspecies of *Ligilactobacillus* (*L. aviarius* subsp. *aviarius* and *L. aviarius* subsp. *araffinosus*) have now been re-classified into different species, namely *L. aviarius* and *L. araffinosus* (1, 17). This split is supported by ANI as well as *pheS* gene analysis, in which the genomes and *pheS* genes of the type strains only share 90% and 94.2% sequence similarity, respectively. The two subspecies of *Latilactobacillus sakei, L. sakei* subsp. *sakei* and *L. sakei* subsp. *carnosus* share only 93.2% *pheS* gene sequence similarity. Similarly, the two subspecies of *Lactiplantibacillus plantarum, L. plantarum* subsp. *plantarum* and *L. plantarum* subsp. *argentoratensis*, share only 91.7% *pheS* gene sequence similarity. However, while the case of *L. plantarum* may warrant further investigation (borderline ANI value between subspecies of 95.7%), ANI analysis of the two *L. sakei* subspecies confirms the standing of *L. sakei* subsp. *carnosus* as a subspecies (ANI: 97.3%).

Both species, *L. plantarum* and *L. sakei*, can be resolved to the subspecies level by our *pheS* taxonomic framework. For the remaining sub-species containing lactobacilli, namely *Loigolactobacillus coryniformis* (subsp. *torquens*), *Lactobacillus delbrueckii* (subsp. *bulgaricus*, subsp. *indicus*, subsp. *jacobsenii*, subsp. *lactis*, and subsp. *sunkii*), *Lactobacillus kefiranofaciens* (subsp. *kefirgranum*), and *Lacticaseibacillus paracasei* (subsp. *tolerans*), all subspecies within the respective species share ≥97% *pheS* gene sequence similarity, supporting their standing in nomenclature as subspecies. These cannot be resolved to the subspecies level by our current *pheS* taxonomic framework. In summary, *pheS*-DB allows differentiation between 263 species-level clades and a further two subspecies (Fig. 2).

The lactobacilli were previously divided into 24 phylogroups based on sequence information of the concatenated protein sequences of single-copy core genes of 174 type strains (6). A later whole genome-based taxonomy of the *Lactobacillus* genus complex represented a total of 208 *Lactobacillus* and *Pediococcus* species all of which clustered within the 24 phylogroups (17). In a recent keystone study that followed a comprehensive polyphasic approach, one additional phylogroup (genus *Acetilactobacillus*) was detected through the inclusion of the latest validly described species, and one phylogroup (*L. salivarius* group) was split into two (1). All 26 phylogroups were then converted into genera with unique genus names (1). In our analysis, a reference sequence for *Acetilactobacillus jinshanensis*, the type species of the genus, is missing. Hence, the 263 species-level clades and 10 subspecies that were included in the *pheS* phylogenetic tree in this study fell into 25 of the 26 genera. All genera formed monophyletic clades apart from three exceptions: 1. The sequence of *Paucilactobacillus kaifaensis* clustered basally to the clade containing the genera *Paucilactobacillus, Limosilactobacillus, Lactiplantibacillus* and *Furfurilactobacillus*, 2. the genus *Schleiferilactobacillus* (former *perolens* phylogroup) formed a cluster within the genus *Lacticaseibacillus* (former *casei* phylogroup), and 3. the genera *Secundilactobacillus* (former *collinoides* phylogroup) and *Ligilactobacillus* (former *salivarius* phylogroup) were paraphyletic, with three species of *Ligilactobacillus* (*L. hayakitensis, L. ruminis*, and *L. salivarius*) clustering into the *Liquorilactobacillus* clade (also former *salivarius* phylogroup; Fig. 2). These difficulties may result from the fact that *pheS* sequences covering the entire amplicon length were not available for all species. The information derived from shorter sequences does not seem to be sufficient in all cases to group the respective species into the correct genus-level clades. This underpins the necessity for whole genome-based approaches to define existing and erect new genera within the lactobacilli. However, it does not hamper the use of the *pheS* gene to assign sequence data to particular species.

### Development of a taxonomic framework (pheS-DB)

Based on our *pheS* gene lactobacilli phylogeny, we compiled a comprehensive *pheS* database (*pheS*-DB). *PheS*-DB comprises a sequence .fasta file of a total of 445 *pheS* gene sequences encompassing 263 species-level clades of lactobacilli (File S1 in the supplemental material) and the corresponding taxonomy .txt file (File S2 in the supplemental material). In addition, we included reference sequences and taxonomic strings of the type strains of the genera *Bacillus, Enterococcus, Fructobacillus, Lactococcus, Leuconostoc, Oenococcus, Streptococcus*, and *Weissella* due to their frequent occurrence in fermented foods and beverages and the likelihood of these species to be captured by the *pheS* primers.

This framework is compatible with commonly used next-generation amplicon sequencing pipelines such as mothur (21) or QIIME2 (23) and can be used for taxonomic assignment of *pheS* gene sequence data. The taxonomy .txt file comprises a total of eight taxonomic levels: domain, phylum, class, order, family, genus, species (in some cases subspecies as described above), and strain (see File S2). We recommend use at the species/subspecies level (pass option “-L 7” with QIIME script “qiime taxa collapse”) for high-resolution lactobacilli community structure analysis.

### Evaluation of detection sensitivity and accuracy of taxonomic assignment through analysis of mock communities

High throughput 16S rRNA and *pheS* gene sequencing of two synthetic (mock) communities allowed us to validate the results obtained with the newly developed lactobacilli framework. For these two samples, after quality filtering, an average of 20,261 ± 11,603 and 32,285 ± 9,016 paired-end sequence reads were obtained for 16S rRNA and *pheS* genes, respectively.

While the 16S rRNA gene allows for coverage of the vast majority of bacterial biodiversity detected to date, non-full length 16S rRNA gene sequences are generally only accurate for taxonomic assignment down to the genus level (25). In our study, based on 16S rRNA genes, only three of a total of 13 species that comprised the mock community were identified correctly at the species level (*Lactiplantibacillus plantarum, Levilactobacillus brevis*, and *Pediococcus acidilactici*). At the genus level, out of the 7 major genera represented in the two samples, five were correctly identified by 16S rRNA gene profiling (*Lacticaseibacillus, Lactiplantibacillus, Lactobacillus, Levilactobacillus*, and *Pediococcus*), while two genera, *Lentilactobacillus* and *Limosilactobacillus*, remained unidentified. *PheS* gene sequence data revealed a considerably higher community structure resolution. All thirteen species included in the artificial community were successfully detected and correctly identified, even those that were spiked into the mock community at levels as low as 0.02 pg per µl DNA. Assuming an average genome size of 2.2 Mbp (3), this detection limit corresponds to ∼10 genome copies per µl template DNA. Closely related species such as *L. plantarum* and *L. pentosus* were successfully distinguished. In the case of *L. plantarum*, the *pheS* methodology even allowed dissecting of the community down to the subspecies level, where *L. plantarum* subsp. *plantarum* and *L. plantarum* subsp. *argentoratensis* were clearly distinguished. For species that were represented in the sample at a relative abundance of ≥0.1%, sequence data showed relative abundances that corresponded well with their expected abundances in the mock community (Fig. 3). Below this threshold, species tended to be overrepresented. This observed deviation from the expected for low template concentrations may have been caused, e.g., by primer bias and preferential amplification of certain DNA templates. As this trend was observed independent of the species, a possible explanation is also that pipette error more severely affected highly diluted DNA of individual lactobacilli species, and thus may have caused consistent overrepresentation of those species that were added into the mock communities at very low abundances. Importantly, primer bias, if present, is expected to similarly affect samples that have been treated in the same way. Under these circumstances, while absolute relative abundances may not be exact, presence or absence, as well as trends in relative abundances of genera and species between different samples (β-diversity) can be measured reliably. In future, mock communities of lactobacilli cells rather than DNAs may provide better insights into the assessment of complex communities and potential biases (e.g., DNA extraction bias) influencing these analyses.

**FIG 3.**
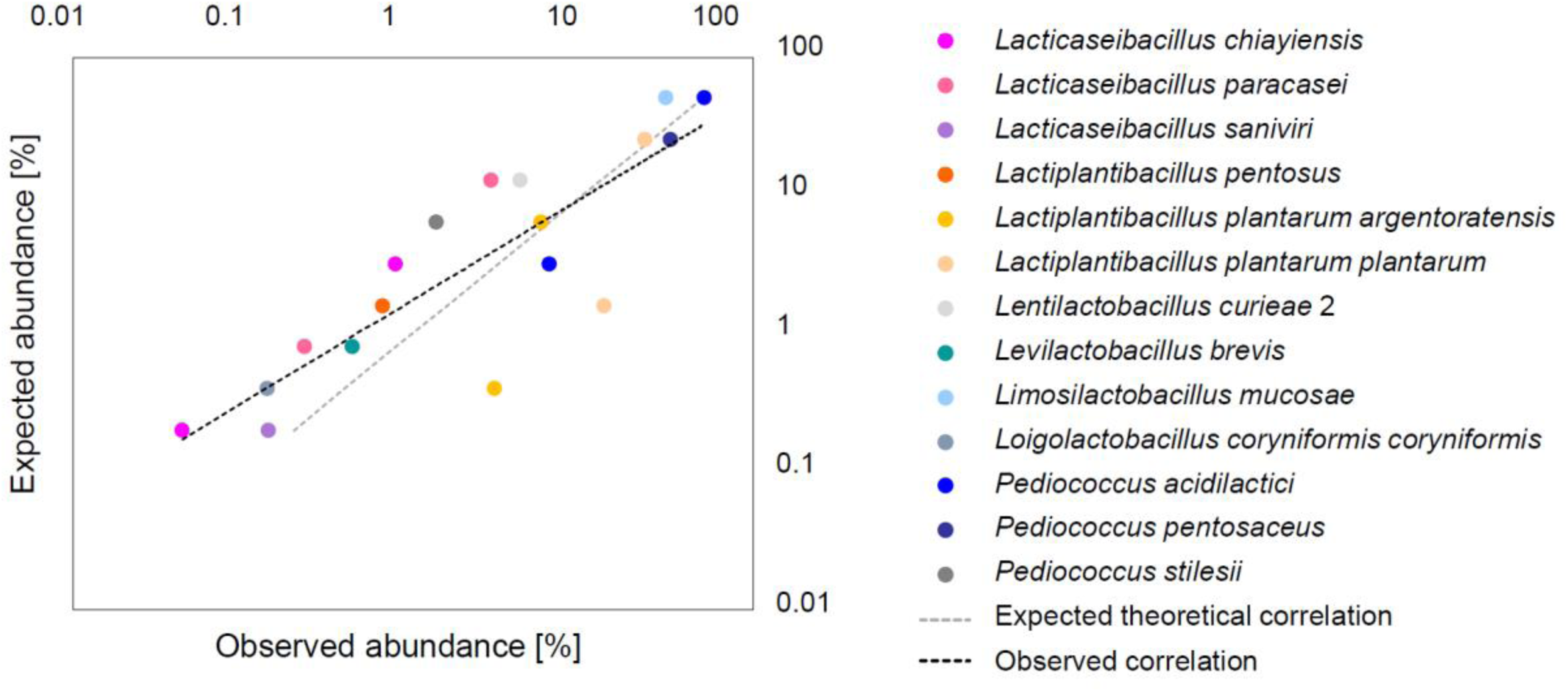
Evaluation of accuracy of the *pheS* gene sequencing and taxonomic assignment approach using two individual lactobacilli mock communities (STD1 and STD2) consisting of DNAs of 13 different species or subspecies, each at different proportions (Tab S2 in the Supplemental Material). The dotted black line indicates the correlation between expected relative abundance and observed relative abundance, while the dotted gray line indicates the theoretically expected perfect correlation. Only species that were represented in either of the mock communities at ≥0.1% relative abundance are shown in the graph.

### Comparative analysis of lactobacilli diversity and community structure in environmental samples based on pheS and 16S rRNA marker genes

To demonstrate the application of our high-resolution *pheS*-based community profiling method, we collected 24 environmental samples from a variety of fermented foods as well as different host-associated matrices. Our aims were, first, to identify samples with largely differing lactobacilli communities for the purpose of isolation, and second, to assess similarities and differences between lactobacilli community structure in similar food matrices. The samples were analyzed by using both 16S rRNA and *pheS* marker gene amplicon sequencing. After filtering, denoising, merging, and chimera removal using DADA2, a total of 723,525 paired-end 16S rRNA gene (30,147 ± 13,339 sequences per sample (average ± standard deviation)) and 810,805 paired-end *pheS* gene sequences were obtained (33,784 ± 11,448 sequences per sample) across the 24 environmental samples (Tab. S3 in the supplemental material). Analysis of 16S rRNA gene sequence data allowed to obtain complete bacterial community profiles from all environmental samples. Members of the lactobacilli accounted for an average of 26.8% ± 15.5% of total 16S rRNA gene sequence reads across all samples. Of these, an average of 56.0% ± 11.2% achieved species-level assignment, however, the accuracy of these species-level assignments may be compromised as discussed above. In the *pheS* approach, lactobacilli made up an average of 55.1% ± 24.3% of total sequence reads across all samples, of which 100% achieved species-level assignment. An average of 35.5% ± 23.8% of *pheS* gene sequence reads remained unassigned (represented by 1,083 representative sequences). Out of these representative sequences, 78.2% could be assigned at least to the genus-level *via* Kraken2 (Fig. S2 in the supplemental material), with the highest percentage of representative sequences observed for the genera *Enterobacter* (14.4%), *Bacillus* (6.1%) and *Lactococcus* (4.4%). Only 45 representative sequences (4.2%) were assigned to the lactobacilli by Kraken2. Upon closer inspection of these sequences by BLAST and alignment, we found that 21 were true *pheS* sequences, while the remaining 24 matched other genomic regions, and likely resulted from unspecific primer binding. Unspecific binding to *pheS* genes of non-target organisms or to other genomic regions of target-or non-target organisms reduces the number of total lactobacilli *pheS* sequence reads per sample and thus needs to be considered. Importantly, however, the 21 representative sequences of true lactobacilli *pheS* gene sequences that remained unassigned with *pheS*-DB, altogether represented sequences that accounted for only 0.2% and 0.07% of unassigned and total sequence reads, respectively. As such, any *pheS*-DB unassigned sequences did not contribute noticeably to the apparent lactobacilli community structure in the environmental samples and were excluded from subsequent analyses in this study.

Results obtained from *pheS* gene profiling were compared to profiles reconstructed *via* partial 16S rRNA gene amplicon sequencing at both genus-(Fig. 4) and species-level (Fig. S1 in the supplemental material). Importantly, only reads classified as belonging to the lactobacilli were included in this analysis. Reads that were classified into a designated species within the lactobacilli but accounted for a relative abundance of <1% in all samples were collapsed as “Lactobacilli <1%”. Reads that were classified to the genus level only (16S rRNA gene data only) were collapsed as “Lactobacilli, other”.

**FIG 4.**
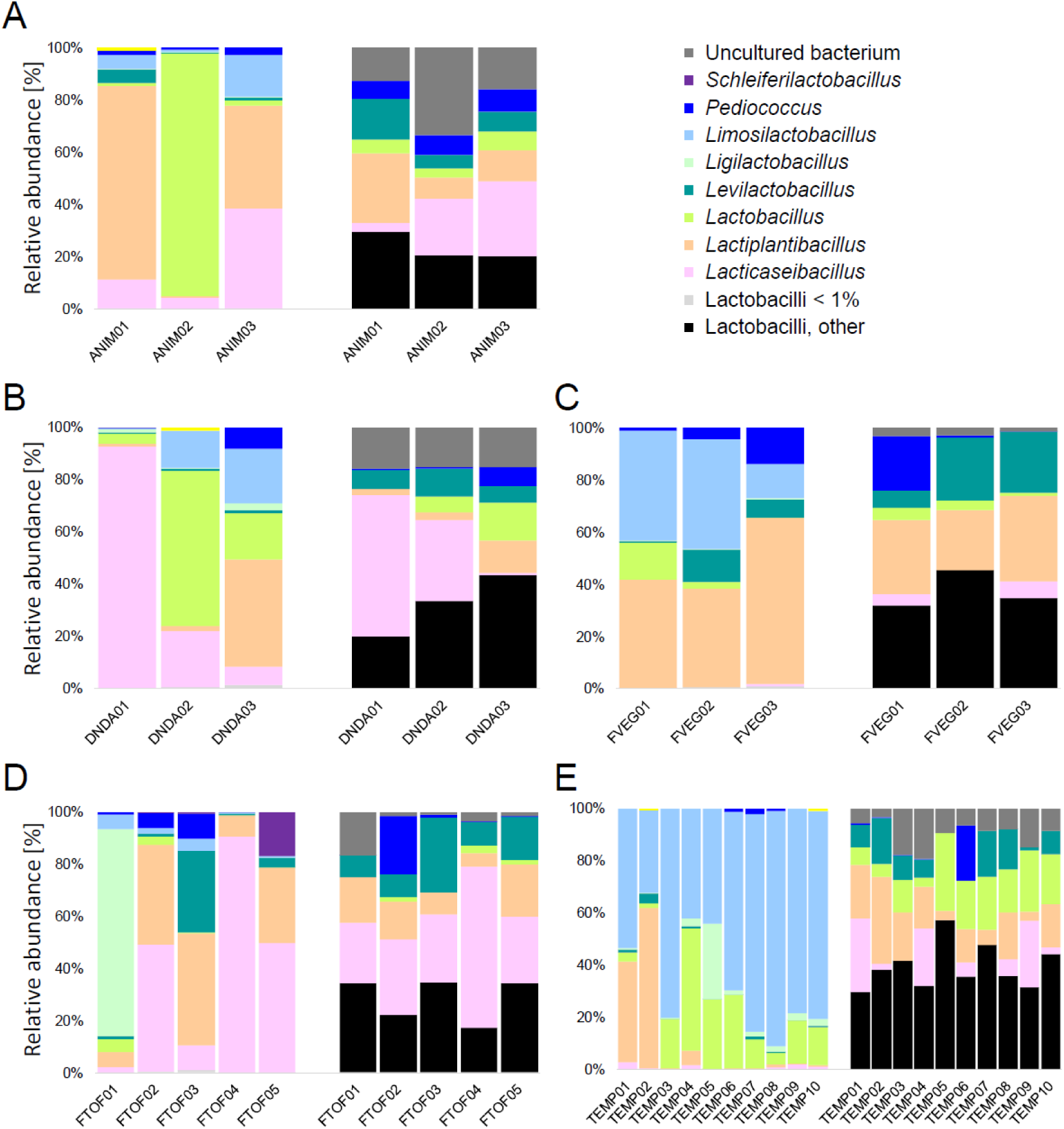
Genus-level community structure of the lactobacilli based on *pheS* (left) and 16S rRNA marker genes (right) in samples from five broad environments. Bar plots show the relative abundances of lactobacilli genera in A. three animal-associated fecal or jejunum samples (ANIM), B. three dairy and non-dairy alternative products (DNDA), C. three fermented vegetable samples (FVEG), D. five fermented tofu samples (FTOF), and E. ten tempeh samples (TEMP). Only genera with a relative abundance of ≥1% in at least one of the 24 samples are reported. Designated genera that showed relative abundances of <1% in all samples are collapsed as “Lactobacilli <1%”, while sequences that could not be assigned to the genus or species level are collapsed as “Lactobacilli, other”.

The *pheS* approach allowed all lactobacilli sequence reads to be assigned down to the species level. Furthermore, the number of detected taxa, that accounted for ≥1% in at least one sample according to the Chao-1 index was significantly higher based on *pheS* than based on 16S rRNA genes (12 *versus* 7 (when taking into account undefined clades), *P*<0.001). Across all analyzed samples, a total of 19 different lactobacilli species (≥1% in at least one sample) were detected by *pheS* gene profiling compared to only five by 16S rRNA gene profiling (Fig. 5). Beta-diversity as calculated with the Bray-Curtis metric showed that lactobacilli communities were generally less similar between samples based on *pheS* gene sequence data (within-method similarity 29.3% ± 23.5%, average ± standard deviation) as compared to 16S rRNA gene sequence data (within-method similarity 63.3% ± 13.8%, average ± standard deviation, *P*<0.001). Between-method similarity was low at only 13.0% ± 10.9% (average ± standard deviation). The differences between methods were corroborated by sample clustering using correlation distance and average linkage (Fig. 5) and may be attributed to i.) a limited number of lactobacilli species that are distinguishable based on the partial 16S rRNA gene as compared to the higher resolving *pheS* gene, and ii.) a lower number of lactobacilli species that are represented by 16S rRNA gene sequences in the SILVA database.

**FIG 5.**
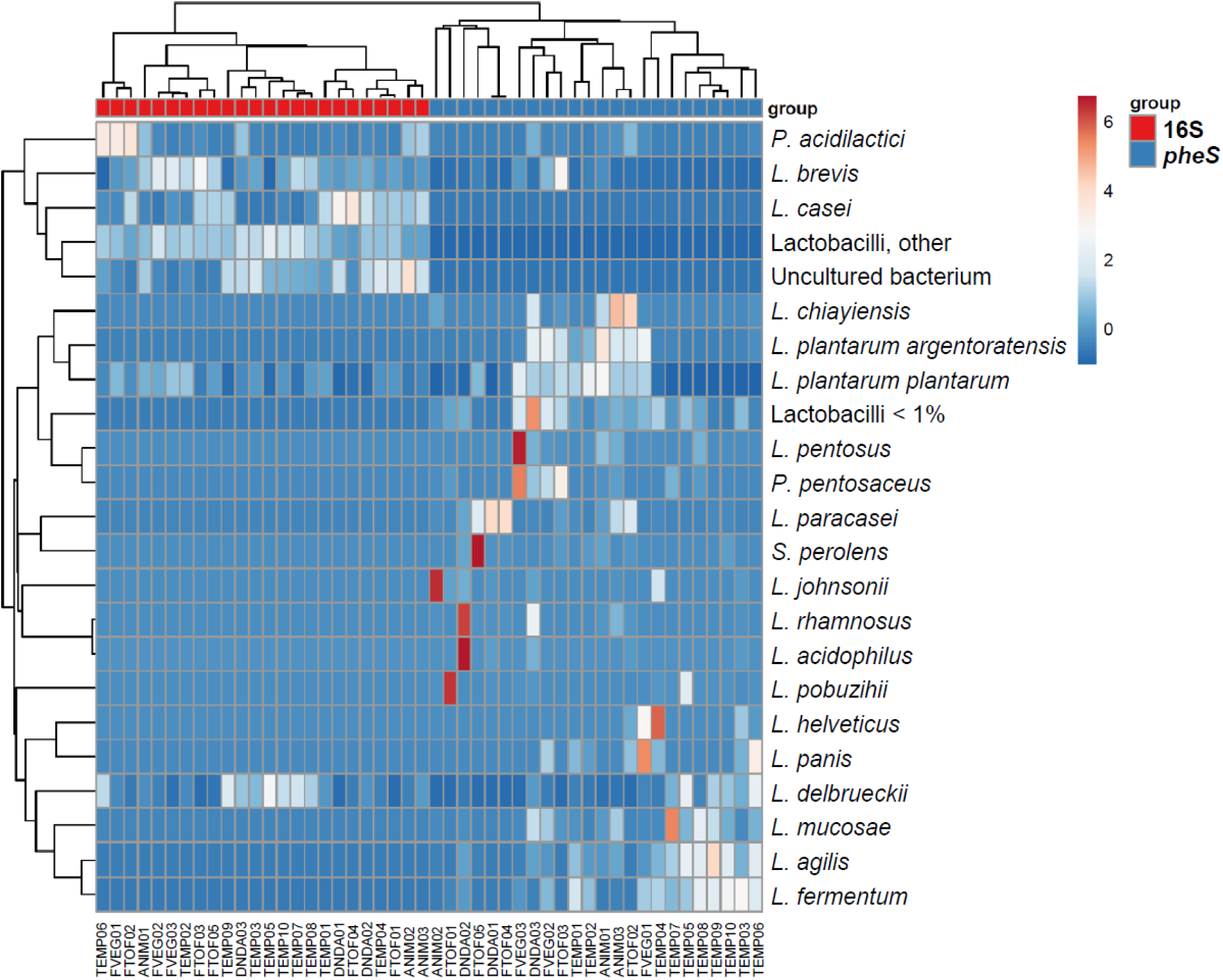
Heatmap and dendrogram representation of within-and between-method sample similarity based on 16S rRNA and *pheS* gene *LSL* group community structure analysis at the species level. Samples are labelled according to Table 1.

The heatmap representation, in particular of the samples analyzed with the *pheS*-based methodology, allowed to identify samples that contained the largest proportions of a particular species. This may be useful for the purpose of isolating additional and potentially novel strains from the analyzed samples (Fig. 5).

**Table 1.** Fermented food and host-associated samples analyzed in this study for bacterial community composition using 16S rRNA and *pheS* gene amplicon sequencing.

| Sample name | Sample type | Sample origin |
| --- | --- | --- |
| DNDA01 | Coconut milk yogurt | Singapore, Singapore |
| DNDA02 | Greek yogurt | Singapore, Singapore |
| DNDA03 | Kefir | Singapore, Singapore |
| ANIM01 | Guinea pig feces | Singapore, Singapore |
| ANIM02 | Mouse jejunum content | Singapore, Singapore |
| ANIM03 | Wild boar feces | Singapore, Singapore |
| FVEG01 | Fermented mustard greens | Ha Long City, Vietnam |
| FVEG02 | Kimchi | Ha Long City, Vietnam |
| FVEG03 | Fermented mustard greens | Jakarta, Indonesia |
| TEMP01 | Tempeh | Pasar Tegal Danas, Cikarang, Jakarta, Indonesia |
| TEMP02 | Tempeh | Lemahebang, Cikarang, Jakarta, Indonesia |
| TEMP03 | Tempeh | Bojong Market, Karawang, Jakarta, Indonesia |
| TEMP04 | Tempeh | Bojong Market, Karawang, Jakarta, Indonesia |
| TEMP05 | Tempeh | Roxy Market, Cikarang, Jakarta, Indonesia |
| TEMP06 | Tempeh | Cikarang Puteh, Cikarang, Jakarta, Indonesia |
| TEMP07 | Tempeh | Rusa Market, Cikarang, Jakarta, Indonesia |
| TEMP08 | Tempeh | Rusa Market, Cikarang, Jakarta, Indonesia |
| TEMP09 | Tempeh | Rusa Market, Cikarang, Jakarta, Indonesia |
| TEMP10 | Tempeh | Niaga Haji Abdul Malik Market, Cikarang, Jakarta, Indonesia |
| FTOF01 | Fermented tofu | Kuala Lumpur, Malaysia |
| FTOF02 | Fermented tofu | Kuala Lumpur, Malaysia |
| FTOF03 | Fermented tofu | Kuala Lumpur, Malaysia |
| FTOF04 | Fermented tofu | Kuala Lumpur, Malaysia |
| FTOF05 | Fermented tofu | Kuala Lumpur, Malaysia |

**Table 2.** Previous species assignment and revised species assignment of several lactobacilli *pheS* gene sequences retrieved from GenBank. For the “previous assignment”, the original genus name “*Lactobacillus*” was kept for clarity, while the re-classified species-level assignments include the new genus names according to Zheng *et al*. (1).

| Accession Number | Previous assignment | Similarity to type strain<br>(previous assignment) | Re-classified assignment | Similarity to type strain<br>(re-classified assignment) |
| --- | --- | --- | --- | --- |
| AM087760 | *Lactobacillus murinus* | 91.3% | *Ligilactobacillus animalis* | 98.3% |
| AM087771<sup>T</sup> | *Lactobacillus homohiochii* | --- | *Fructilactobacillus fructivorans* | 100% |
| AM284237 | *Lactobacillus gasseri* | 95.4% | *Lactobacillus paragasseri* | 100% |
| AM284241 | *Lactobacillus gasseri* | 95.4% | *Lactobacillus paragasseri* | 100% |
| AM284196 | *Lactobacillus gasseri* | 95.6% | *Lactobacillus paragasseri* | 99.3% |
| AM284194 | *Lactobacillus gasseri* | 95.4% | *Lactobacillus paragasseri* | 99.1% |
| AM284246 | *Lactobacillus farciminis* | 93.6% | *Companilactobacillus formosensis* | 99.1% |

**Table 3.** Sequence similarities based on *pheS* genes (*pheS*-based) and whole genomes (ANI-based) between closely related strains of different species or distantly related strains within the same species. Clade names refer to the species-level (L7) assignment as provided in *pheS*-DB. The same clade name is given if type strains of the species cannot be confidently distinguished based on *pheS* genes. Species-level clades are divided into subclades, if strains of the same species are only distantly related to the type strain (*pheS* gene sequence similarity <90%, ANI <95%). For the two *C. crustorum* strains, only the type strain genome is publicly available, and no comparison can be done at the genomic level.

| (Type) strain(s) present in clade | Given clade name | Sequence similarity between strains |  |
| --- | --- | --- | --- |
|  |  | *pheS*-based | ANI-based |
| *A. kosoi* NBRC 113063<sup>T</sup> | *A. kosoi/micheneri/timberlakei* |  |  |
| *A. micheneri* DSM 104126<sup>T</sup> | *A. kosoi/micheneri/timberlakei* | 98.4–99.5% | 94.3–97.6% |
| *A. timberlakei* DSM 104128<sup>T</sup> | *A. kosoi/micheneri/timberlakei* |  |  |
| *C. mishanensis* LMG 31048<sup>T</sup> | *C. mishanensis/salsicarnum* | 100% | 99.9% |
| *C. salsicarnum* LMG 31401<sup>T</sup> | *C. mishanensis/salsicarnum* |  |  |
| *L. amylovorus* DSM 20531<sup>T</sup> | *L. amylovorus/kitasatonis* | 98.5% | 92.4% |
| *L. kitasatonis* JCM 1039<sup>T</sup> | *L. amylovorus/kitasatonis* |  |  |
| *C. crustorum* LMG 23699<sup>T</sup> | *C. crustorum* | 89.9% | Only type strain sequence |
| *C. crustorum* LMG 23702 | *C. crustorum* 2 |  | available |
| *L. curieae* CCTCC M 2011381<sup>T</sup> | *L. curieae* | 84.4% | 83.8% |
| *L. curieae* NBRC111893 | *L. curieae* 2 |  |  |

Furthermore, Principal Component Analysis revealed a strong influence of sample type on clustering of TEMP samples, with FVEG samples also grouping relatively closely together (Fig. 6). Interestingly, the remaining sample types, ANIM, DNDA, and FTOF, were distributed across two clusters (Fig. 5, 6). Cluster 1 was characterized by the prevalence of *L. plantarum* subsp. *plantarum* and included two fermented tofu samples, one kefir sample, one wild boar and one guinea pig fecal sample. Samples in cluster 2 included three fermented tofu samples, one mouse jejunum content sample, one coconut milk and one Greek yogurt sample. This cluster was characterized by the presence of *L. paracasei* in all six samples and relatively high abundances of *L. pobuzihii, L. johnsonii*, or *L. acidophilus*. To some degree, these samples shared similar lactobacilli communities, however, the broad clustering observed here may be a result of the small sample size per sample type.

**FIG 6.**
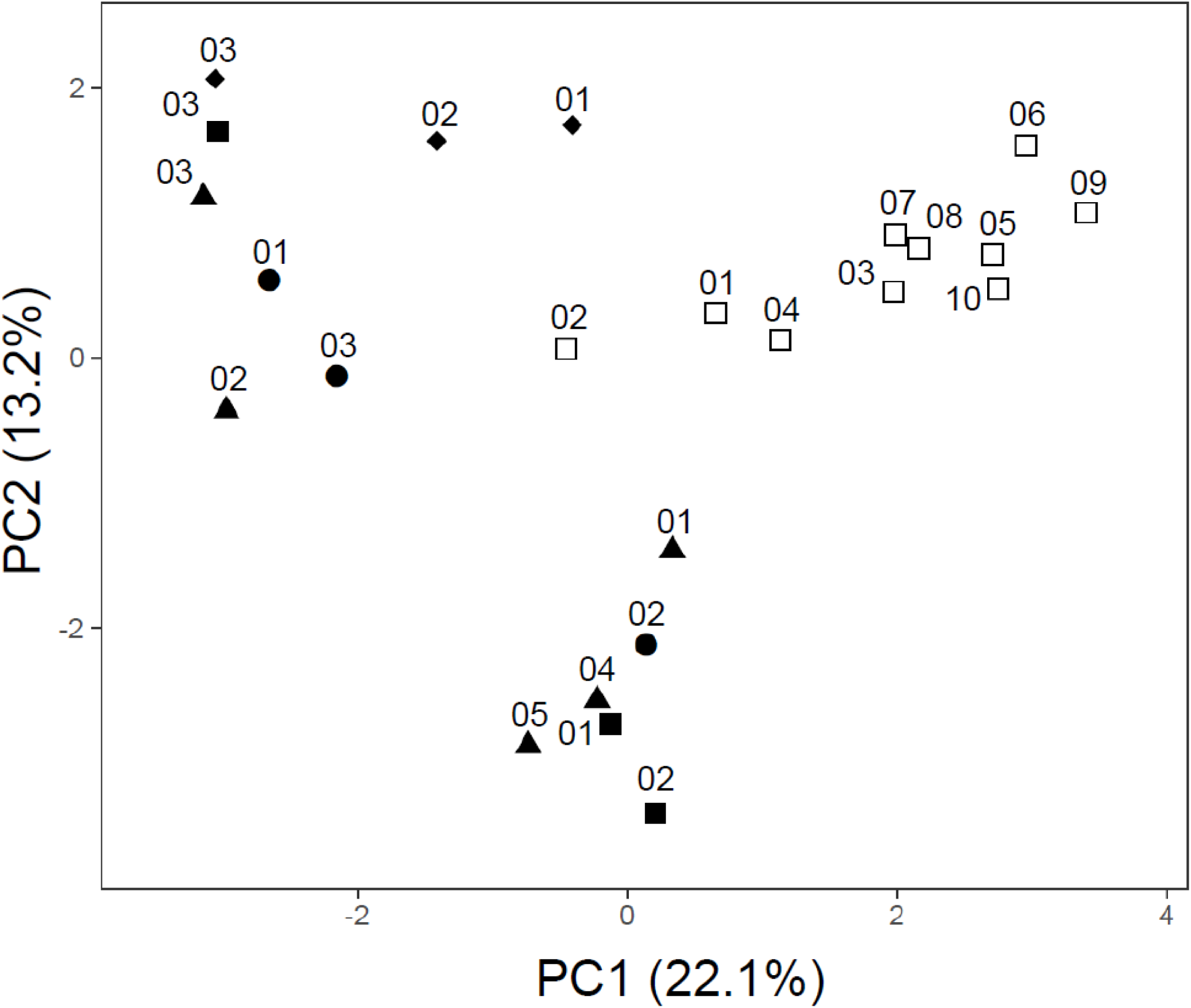
Principal Component Analysis of lactobacilli community structure in all analyzed samples based on *pheS* genes. Sample identifiers refer to Table 1 with symbols representing the five different sample types as follows: ANIM (•), DNDA (▪), FVEG (♦s), TEMP (□), FTOF (▴).

Taken together, the *pheS* approach allowed to confirm populations that had previously been reported from specific sample types, as well as to identify populations of lower but physiologically relevant relative abundances that had not been linked to particular intestinal environments or food fermentation processes previously.

With third generation technology becoming more available and affordable, high-throughput sequencing of full-length 16S rRNA genes may soon be achievable routinely (25). However, sophisticated denoising algorithms are required to correct for PCR and sequencing errors. Moreover, in the case of some species of lactobacilli, even full-length 16S rRNA gene sequencing may not be able to resolve sequences at the species-level (52). The *pheS* approach may be improved in future by extending primer coverage and by obtaining *pheS* gene sequences for the remaining six type strains, since unavailability of these missing species currently renders them elusive to detection and relative quantification in environmental samples. Despite these limitations, our new *pheS* gene sequencing approach combined with taxonomic assignment through *pheS*-DB, provides highly accurate and resolved species-level reconstruction of lactobacilli communities and may complement 16S rRNA gene profiling of a broad range of complex environmental and host-associated matrices.

## CONCLUSIONS

The newly established *pheS* gene sequencing approach together with taxonomic assignment by *pheS*-DB provides comprehensive and highly resolving community structure information of lactobacilli. The framework currently enables exact species-level taxonomic assignment of a total of 263 lactobacilli species and a further two subspecies (*L. plantarum* subsp. *argentoratensis*, and *L. sakei* subsp. *carnosus*). The *pheS* gene amplicon sequencing approach is highly sensitive with a limit of detection of approximately 0.02 pg per μl DNA (corresponding to ∼10 genome copies). We applied the method to 24 diverse samples stemming from host-associated environments, dairy and non-dairy alternative products, fermented Asian soybean products (tempeh and tofu), as well as fermented vegetables and were able to evaluate diversity and community composition of the resident lactobacilli at great depth. As such, the newly described methodology represents a valuable tool for understanding lactobacilli diversity in a broad range of sample types. In future, it may also be used to monitor lactobacilli communities during industrial fermentation processes, where it could potentially assist in linking distinct populations to desirable or non-desirable food or feed rheological and organoleptic properties or metabolites.

## ACKNOWLEDGMENTS

This study was funded by Wilmar International Limited (WIL@NUS Corporate Laboratory, Singapore). We thank our collaborators Benjamin Lee, Bryan Lim, and Chanelle Lim (all NParks, Singapore) for assistance in the collection of a wild boar fecal sample, as well as Antonio Suwanto, Esti Puspitasari (both Wilmar Indonesia), Peh Ping-Teik and Do Thi Kim Oanh (both Wilmar CLV) for help with the collection of fermented food samples in Indonesia and Vietnam. We are indebted to Connie Chew Sim Hwa (Kuala Lumpur, Malaysia) for allowing us to collect samples from her home-fermented tofu.

## REFERENCES

1. Zheng J, Wittouck S, Salvetti E, Franz CM, Harris HM, Mattarelli P, O’Toole PW, Pot B, Vandamme P, Walter J, Watanabe K et al. 2020. A taxonomic note on the genus Lactobacillus: Description of 23 novel genera, emended description of the genus Lactobacillus Beijerinck 1901, and union of Lactobacillaceae and Leuconostocaceae. Int J Syst Evol Micr. 10.1099/ijsem.0.004107.

2. Douillard FP, De Vos WM. 2014. Functional genomics of lactic acid bacteria: from food to health. Microb Cell Fact 13:S1–S8.

3. Duar RM, Lin XB, Zheng J, Martino ME, Grenier T, Pérez-Muñoz ME, Leulier F, Gänzle M, Walter J. 2017. Lifestyles in transition: evolution and natural history of the genus Lactobacillus. FEMS Microbiol Rev 41:S27–S48.

4. Duar RM, Frese SA, Lin XB, Fernando SC, Burkey TE, Tasseva G, Peterson DA, Blom J, Wenzel CQ, Szymanski CM, Walter J. 2017. Experimental evaluation of host adaptation of Lactobacillus reuteri to different vertebrate species. Appl Environ Microbiol 83:e00132–17.

5. Sun Z, Harris HM, McCann A, Guo C, Argimón S, Zhang W, Yang X, Jeffery IB, Cooney JC, Kagawa TF, Liu W, et al. 2015. Expanding the biotechnology potential of lactobacilli through comparative genomics of 213 strains and associated genera. Nat Commun 6:1–3.

6. Zheng J, Ruan L, Sun M, Gänzle M. 2015. A genomic view of lactobacilli and pediococci demonstrates that phylogeny matches ecology and physiology. Appl Environ Microbiol 81:7233–7243.

7. Bokulich NA, Mills DA. 2012. Next-generation approaches to the microbial ecology of food fermentations. BMB Rep 45:377–389.

8. Gänzle MG. 2015. Lactic metabolism revisited: metabolism of lactic acid bacteria in food fermentations and food spoilage. Curr Opin Food Sci 2:106–117.

9. Staley JT, Konopka A. 1985. Measurements of in situ activities of non-photosynthetic microorganisms in aquatic and terrestrial habitats. Annu Rev Microbiol 39:321–346.

10. De Filippis F, Parente E, Ercolini D. 2017. Metagenomics insights into food fermentations. Microb Biotechnol 10:91–102.

11. Agyirifo DS, Wamalwa M, Otwe EP, Galyuon I, Runo S, Takrama J, Ngeranwa J. 2019. Metagenomics analysis of cocoa bean fermentation microbiome identifying species diversity and putative functional capabilities. Heliyon 5:e02170.

12. Verce M, De Vuyst L, Weckx S. 2019. Shotgun metagenomics of a water kefir fermentation ecosystem reveals a novel Oenococcus species. Front Microbiol 10:479.

13. Scholz M, Ward DV, Pasolli E, Tolio T, Zolfo M, Asnicar F, Truong DT, Tett A, Morrow AL, Segata N. Strain-level microbial epidemiology and population genomics from shotgun metagenomics. Nat Methods 13:435–438.

14. Schloss PD. 2020. Reintroducing mothur: 10 years later. Appl Environ Microbiol 86:e02343–19.

15. Parks DH, Chuvochina M, Waite DW, Rinke C, Skarshewski A, Chaumeil PA, Hugenholtz P. 2018. A standardized bacterial taxonomy based on genome phylogeny substantially revises the tree of life. Nat Biotechnol 36:996–1004.

16. Salvetti E, Harris HM, Felis GE, O’Toole PW. 2018. Comparative genomics of the genus Lactobacillus reveals robust phylogroups that provide the basis for reclassification. Appl Environ Microbiol 84:e00993–18.

17. Wittouck S, Wuyts S, Meehan CJ, van Noort V, Lebeer S. 2019. A genome-based species taxonomy of the Lactobacillus Genus Complex. mSystems 4:e00264–19.

18. McDonald D, Price MN, Goodrich J, Nawrocki EP, DeSantis TZ, Probst A, Andersen GL, Knight R, Hugenholtz P. 2012. An improved Greengenes taxonomy with explicit ranks for ecological and evolutionary analyses of bacteria and archaea. ISME J 6:610–618.

19. Quast C, Pruesse E, Yilmaz P, Gerken J, Schweer T, Yarza P, Peplies J, Glöckner FO. 2013. The SILVA ribosomal RNA gene database project: improved data processing and web-based tools. Opens external link in new window. Nucleic Acids Res 41:D590–D596.

20. Yilmaz P, Parfrey LW, Yarza P, Gerken J, Pruesse E, Quast C, Schweer T, Peplies J, Ludwig W, Glöckner FO. 2014. The SILVA and “all-species living tree project (LTP)” taxonomic frameworks. Nucleic Acids Res 42:D643–648.

21. Schloss PD, Westcott SL, Ryabin T, Hall JR, Hartmann M, Hollister EB, Lesniewski RA, Oakley BB, Parks DH, Robinson CJ, Sahl JW. 2009. Introducing mothur: open-source, platform-independent, community-supported software for describing and comparing microbial communities. Appl Environ Microbiol 75:7537–7541.

22. Caporaso JG, Kuczynski J, Stombaugh J, Bittinger K, Bushman FD, Costello EK, Fierer N, Pena AG, Goodrich JK, Gordon JI, Huttley GA, Kelley ST, Knights D, Koenig JE, Ley RE, Lozupone CA, McDonald D, Muegge BD, Pirrung M, Reeder J, Sevinsky JR, Turnbaugh PJ, Walters WA, Widman J, Yatsunenko T, Zaneveld J, Knight R. 2010. QIIME allows analysis of high-throughput community sequencing data. Nat Methods 7:335–336.

23. Bolyen E, Rideout JR, Dillon MR, Bokulich NA, Abnet CC, Al-Ghalith GA, Alexander H, Alm EJ, Arumugam M, Asnicar F, Bai Y, et al. 2019. Reproducible, interactive, scalable and extensible microbiome data science using QIIME 2. Nature Biotechnol 37:852–857.

24. Stackebrandt E, Ebers J. 2006. Taxonomic parameters revisited: tarnished gold standards. Microbiol Today 33:152–155.

25. Johnson JS, Spakowicz DJ, Hong BY, Petersen LM, Demkowicz P, Chen L, Leopold SR, Hanson BM, Agresta HO, Gerstein M, Sodergren E. 2019. Evaluation of 16S rRNA gene sequencing for species and strain-level microbiome analysis. Nat Commun 10:1–1.

26. Naser SM, Thompson FL, Hoste B, Gevers D, Dawyndt P, Vancanneyt M, Swings J. 2005. Application of multilocus sequence analysis (MLSA) for rapid identification of Enterococcus species based on rpoA and pheS genes. Microbiology 151:2141–2150.

27. Nyanzi R, Jooste PJ, Cameron M, Witthuhn C. 2013. Comparison of rpoA and pheS gene sequencing to 16S rRNA gene sequencing in identification and phylogenetic analysis of LAB from probiotic food products and supplements. Food Biotechnol 27:303–327.

28. De Filippis F, La Storia A, Stellato G, Gatti M, Ercolini D. 2014. A selected core microbiome drives the early stages of three popular Italian cheese manufactures. PLOS ONE 9:e89680.

29. Parente E, Guidone A, Matera A, De Filippis F, Mauriello G, Ricciardi A. 2016. Microbial community dynamics in thermophilic undefined milk starter cultures. Int J Food Microbiol 217:59–67.

30. Ricciardi A, De Filippis F, Zotta T, Facchiano A, Ercolini D, Parente E. 2016. Polymorphism of the phosphoserine phosphatase gene in Streptococcus thermophilus and its potential use for typing and monitoring of population diversity. Int J Food Microbiol 236:138–147.

31. Milani C, Duranti S, Mangifesta M, Lugli GA, Turroni F, Mancabelli L, Viappiani A, Anzalone R, Alessandri G, Ossiprandi MC, van Sinderen D. 2018. Phylotype-level profiling of lactobacilli in highly complex environments by means of an internal transcribed spacer-based metagenomic approach. Appl Environ Microbiol 84:e00706–18.

32. Xie M, Pan M, Jiang Y, Liu X, Lu W, Zhao J, Zhang H, Chen W. 2019. groEL gene-based phylogenetic analysis of Lactobacillus species by high-throughput sequencing. Genes 10:530.

33. Naser SM, Dawyndt P, Hoste B, Gevers D, Vandemeulebroecke K, Cleenwerck I, Vancanneyt M, Swings J. 2007. Identification of lactobacilli by pheS and rpoA gene sequence analyses. Int J Syst Evol Micr 57:2777–2789.

34. Edgar RC. 2004. MUSCLE: multiple sequence alignment with high accuracy and high throughput. Nucleic Acids Res 32:1792–1797.

35. Crooks GE, Hon G, Chandonia JM, Brenner SE. 2004. WebLogo: a sequence logo generator. Genome Res 14:1188–1190.

36. Martin M. 2011. Cutadapt removes adapter sequences from high-throughput sequencing reads. EMBnet. J 17:10–12.

37. Ludwig W, Strunk O, Westram R, Richter L, Meier H Yadhukumar, Buchner A, Lai T, Steppi S, Jobb G, Förster W. 2004. ARB: a software environment for sequence data. Nucleic Acids Res 32:1363–1371.

38. Pritchard L, Glover RH, Humphris S, Elphinstone JG, Toth IK. 2016. Genomics and taxonomy in diagnostics for food security: soft-rotting enterobacterial plant pathogens. Anal Methods-UK 8:12–24.

39. Richter M, Rosselló-Móra R. 2009. Shifting the genomic gold standard for the prokaryotic species definition. P Natl Acad Sci USA 106:19126–19131.

40. Rius AG, Kittelmann S, Macdonald KA, Waghorn GC, Janssen PH, Sikkema E. 2012. Nitrogen metabolism and rumen microbial enumeration in lactating cows with divergent residual feed intake fed high-digestibility pasture. J Dairy Sci 95:5024–5034.

41. Parada AE, Needham DM, Fuhrman JA. 2016. Every base matters: assessing small subunit rRNA primers for marine microbiomes with mock communities, time series and global field samples. Environ Microbiol 18:1403–1414.

42. Apprill A, McNally S, Parsons R, Weber L. 2015. Minor revision to V4 region SSU rRNA 806R gene primer greatly increases detection of SAR11 bacterioplankton. Aquat Microb Ecol 75:129–137.

43. Thompson LR, Sanders JG, McDonald D, Amir A, Ladau J, Locey KJ, Prill RJ, Tripathi A, Gibbons SM, Ackermann G, Navas-Molina JA. 2017. A communal catalogue reveals Earth’s multiscale microbial diversity. Nature 551:457–463.

44. Callahan BJ, McMurdie PJ, Rosen MJ, Han AW, Johnson AJ, Holmes SP. 2016. DADA2: high-resolution sample inference from Illumina amplicon data. Nat Methods 13:581–3.

45. Rognes T, Flouri T, Nichols B, Quince C, and Mahé F. 2016. Vsearch: a versatile open source tool for metagenomics. PeerJ 4:e2584.

46. Wood DE, Lu J, Langmead B. 2019. Improved metagenomic analysis with Kraken 2. Genome Biol 20:257.

47. Hammer Ø, Harper DA, Ryan PD. 2001. PAST: Paleontological statistics software package for education and data analysis. Palaeontol Electron 4:9.

48. Bray JR, Curtis JT. 1957. An ordination of the upland forest communities of Southern Wisconsin. Ecol Monogr 27:325–349.

49. Metsalu T, Vilo J. 2015. ClustVis: a web tool for visualizing clustering of multivariate data using Principal Component Analysis and heatmap. Nucleic Acids Res 43:W566–570.

50. Lei X, Sun G, Xie J, Wei D. 2013. Lactobacillus curieae sp. nov., isolated from stinky tofu brine. Int J Syst Evol Microbiol 63:2501–2505.

51. Scheirlinck I, Van der Meulen R, Van Schoor A, Huys G, Vandamme P, De Vuyst L, Vancanneyt M. 2007. Lactobacillus crustorum sp. nov., isolated from two traditional Belgian wheat sourdoughs. Int J Syst Evol Micr 57:1461–1467.

52. Collins MD, Rodrigues U, Ash C, Aguirre M, Farrow JA, Martinez-Murcia A, Phillips BA, Williams AM, Wallbanks S. 1991. Phylogenetic analysis of the genus Lactobacillus and related lactic acid bacteria as determined by reverse transcriptase sequencing of 16S rRNA. FEMS Microbiol Lett 77:5–12.

